# StabilizeIT: An Automated Workflow for Protein Stabilization

**DOI:** 10.1101/2025.10.09.681370

**Authors:** Nitzan Kutnowski, Yair Budic, Michael Chalick, Noga Alon, Itay Levin, Gideon Lapidoth, Lior Zimmerman

## Abstract

The industrial application of enzymes is often hampered by poor stability and low expression yields. While computational tools can predict stabilizing mutations, many are bound by restrictive licenses that hinder their broader adoption. To address this, we developed **StabilizeIT**, a powerful, open-access webserver for enhancing protein stability and expression.

StabilizeIT integrates a pipeline of curated open-source tools such as ProteinMPNN, AlphaFold2 and SaProt with our state-of-the-art model, SolvIT, which accurately predicts heterologous expression titers in *E. coli*. This unique combination allows for the simultaneous optimization of melting temperature (Tm) and solubility. The pipeline exhibits remarkable speed, generating dozens of high-quality candidates with predicted high titers and increased stability in under an hour, streamlining the path to experimental validation.

To demonstrate its efficacy, StabilizeIT was used to engineer multiple enzymes in our novel biosynthetic pathway for Hyaluronic Acid. The resulting variants showed greatly enhanced thermal stability and expression, proving the pipeline’s real-world utility. StabilizeIT is now available to the community, offering an accessible and validated solution to accelerate the development of robust proteins for diverse applications.

The webserver is freely available at https://stabilizeit.enzymit.com

## Introduction

Engineering proteins for improved stability is a longstanding challenge in biotechnology and medicine. Natural enzymes and therapeutic proteins are often optimized by evolution for function rather than stability^1^. As a result, many wild-type proteins suffer from poor solubility, low thermal stability, or aggregation, limiting their industrial and therapeutic use^2^. Common lab techniques, such as Directed Evolution (DE), require extensive mutagenesis and screening, which is both labor-intensive and resource-demanding^3^. Computational approaches have emerged to reduce this experimental burden by predicting beneficial mutations in silico^4^. Protein stabilization algorithms like PROSS (Protein Repair One-Stop Shop) combine evolutionary sequence information with Rosetta energy calculations to design mutations that enhance protein expression and thermostability^5^. Similarly, web servers such as FireProt^6^ and HotSpot Wizard^7^ automated the identification of stabilizing substitutions using consensus sequences, phylogenetic analysis, and physics-based scoring, often yielding multi-point mutants that significantly raise melting temperatures. However, these methods rely mainly on evolutionary sequence information to guide mutation selection (so conserved amino acids have a higher tendency to get selected as beneficial mutations) and lack higher order context awareness that could affect protein folding, and by extension protein stability

Recent advances in deep learning have opened new avenues for protein design. Large protein language models such as Meta’s ESM2 and SaProt^8^ which are trained on billions of amino acid sequences can assess the “likelihood” of sequences and when trained with supervision, can predict various properties like cellular localization, solubility, thermal stability and even predict 3D structures. Similarly, protei n inverse-folding models such as ProteinMPNN, a deep graph neural network, can generate novel sequences predicted to fold into a given 3D backbone structure^9^. ProteinMPNN designs have shown remarkably high stability and fold accuracy when applied to novel protein scaffolds. Often, redesigned native proteins are predicted to fold more confidently than the wild-types^10^, which suggest an improved stability and decreased unfolding tendencies However, since enzymes require precise structural-functional accuracy for their catalytic activity, when used without structural or functional constraints, ProteinMPNN often produces protein sequences that are catalytically inferior to the wildtype enzyme template^10^. Therefore, to make ProteinMPNN a reliable protein optimization platform, we must layer in knowledge about function, binding sites, and sequence motifs. This transforms the model from a fold-only designer into a comprehensive optimization toolbox for stability and activity.^11,12^

Here we present StabilizeIT - an automated pipeline that integrates state-of-the-art components to stabilize proteins in a reproducible, high-throughput manner. The goal of StabilizeIT is to generate a focused shortlist of variants that maintain functionality, foldability, solubility, and exhibit improved thermal stability. To achieve this, StabilizeIT combines five key modules in a single workflow: **(1)** evolutionary analysis to identify and preserve conserved or critical residues, **(2)** deep generative sequence design with ProteinMPNN, **(3)** sequence and structure-based triaging using language model scores and AlphaFold2 predictions, **(4)** biophysical property prediction (stability ΔΔG and solubility), and **(5)** aggregation of all metrics for easy ranking. By automating these steps with Snakemake^13^ and containerization, the pipeline ensures reproducibility and scalability across computing environments. It is available via a web interface at https://stabilize.enzymit.com. In the following sections, we describe the StabilizeIT workflow in detail and demonstrate its performance on several enzyme templates.

## Methods

### Computational

#### Input and Conservation Analysis

The pipeline accepts either a Protein Data Bank structure file (.pdb) or an amino acid sequence in FASTA format as input. In case a sequence is provided, the structure is predicted using AlphaFold2^14^. From the amino acid sequence the first step is a position-wise analysis of evolutionary conservation, this step is conducted to identify positions that should be kept fixed during design. This step includes a multiple sequence alignment (MSA) of the input sequence against a large protein database UniRef30^15^ using MMseqs2^16^, a fast sequence search tool. From the MSA, a position-specific scoring matrix (PSSM) and conservation frequencies are calculated. By default, the pipeline follows the logic described in *Sumida et-al*^10^ in which residues with ≥50% sequence conservation are considered functionally or structurally important and are fixed (not allowed to change) in subsequent design steps. This ensures that active-site residues and other critical motifs remain unchanged, focusing the design on less-conserved regions.

#### Deep Sequence Design with ProteinMPNN

StabilizeIT uses ProteinMPNN to generate a large diversity of mutated sequences for the target protein. ProteinMPNN takes as an input the backbone coordinates of the query protein (either the provided structure or a predicted structure for sequence-only inputs) and a mask of fixed positions, both the one supplied by the user and the one that was calculated based on conservation. It then proposes new amino acid identities for the non-fixed positions, effectively exploring the sequence space while respecting the backbone geometry and any conserved residues. In this pipeline, the *soluble* model variant^17^ of ProteinMPNN is used. By default, N = 100,000 sequences are sampled for each protein (this number is user-configurable). The output is a FASTA file of all designed sequences. Notably, these designs often contain multiple substitutions relative to the wild type; typically mutating 80%-70% of the non-fixed residues, a scope that would be impractical to search via single-point mutation scanning alone. The assumption is that some of these globally optimized sequences will adopt the target fold with greater stability, as suggested by ProteinMPNN’s success in other design applications.

#### Initial Triage by ESM Inverse Folding

Evaluating the 100 000 candidate sequences generated by the previous ProteinMPNN step experimentally or via full structure prediction is infeasible, so StabilizeIT begins with a rapid in silico triage using Meta’s ESM-IF1^12^ inverse-folding model. For each design, ESM-IF1 evaluates how well the amino-acid sequence conforms to the template backbone, prioritizing those with a high likelihood of correctly adopting the input structure. Because maintaining the precise fold is essential for preserving function while improving other properties, this step filters out sequences unlikely to fold as intended. Importantly, ESM-IF1 operates independently of the ProteinMPNN generator - providing an orthogonal assessment of sequence-structure compatibility before any downstream optimization. Subsequently the pipeline sorts all designs by the ESM-IF1 score and selects the top K sequences for detailed structural evaluation. By default, K = 100 (i.e. 0.1% of designs) proceed to the next stage, though this can be adjusted based on desired thoroughness and computational resources. The output of this step is a shortlist FASTA containing the highest-scoring variants.

#### Structure Prediction with AlphaFold2/ColabFold

The top K sequences are next subjected to 3D structure prediction to evaluate their folding and stability. StabilizeIT employs AlphaFold2 (via the ColabFold toolkit) to predict a structure for each designed variant. ColabFold enables accelerated AlphaFold inference by using a fast MSA generation step with MMseqs2 and reusing one MSA for all generated sequences, making it practical to model 100 proteins in a reasonable time. The result is a set of PDB files, each for a designed variant along with JSON files containing per-residue confidence metrics (pLDDT scores). All these predictions take advantage of GPU acceleration; for example, on an NVIDIA RTX A6000, predicting 100 structures (one target with K=100) takes on the order of 1–2 hours. In summary, the structural modeling step provides atomic-level models for each candidate, enabling the next stage of detailed analysis.

#### Computing Stability and Design Metrics

After structure prediction, StabilizeIT annotates each designed variant with a rich set of metrics to quantify its quality and predicted stability. These computations are orchestrated by a sub-workflow that collates inputs from several tools:

- **Structure deviation (RMSD):** The pipeline calculates the backbone atom’s root-mean-square deviation of each designed protein’s predicted structure from the wild-type structure. Both the global RMSD and an “active-site RMSD” (if an active site or binding region is defined by the user) are computed to ensure the design has not significantly drifted in overall fold or at key functional positions.
- **Confidence scores (pLDDT):** For each design, the mean pLDDT (predicted Local Distance Difference Test) from AlphaFold2 is reported as a measure of model confidence^18^. High pLDDT (close to 90–95) suggests the design sequence is predicted to fold robustly into a well-defined structure, whereas low pLDDT in some regions might flag unstable or disordered segments. The pipeline records both the overall average pLDDT and the average pLDDT on any specified active-site residues which have a particular significance when evaluating enzymes.
- **Sequence identity:** The percentage of identical residues between the designed sequence and the wild-type is calculated. (e.g. a design with 95% identity has only a few mutations, whereas one with 60% identity is heavily mutated). Often, higher mutational density is acceptable if the model confidence remains high, but this metric provides a good risk approximation.
- **ESM and MPNN scores:** same scores that were obtained in the triage step, those scores are attached to the final metrics table. MPNN scores are obtained from the same model used to generate the sequences.
- **Predicted ΔΔG / Tm (thermal stability):** To estimate changes in stability, we integrate the tool SaProt^8^, a recently developed protein language model that predicts protein thermostability changes. SaProt uses a structure-aware transformer model trained on protein melting temperature (Tm) data, providing a proxy for Tm for a given mutation or redesigned sequence. For each design, the pipeline runs SaProt’s pretrained model to predict a relative stability score or Tm. A positive Tm prediction suggests the variant is likely more stable (higher melting point) than the original protein. SaProt predictions, while computational, have shown improved correlation with experimental stability changes by incorporating structural information^19^
- **Predicted solubility:** Similarly, we assess each variant’s solubility using SolvIT, an ensemble of graph neural network models that predict soluble expression in *E. coli*^20^. SolvIT was recently introduced as a means to prioritize designs that are more likely to express well in bacterial systems by outputting a solubility score. SolvIT thus provides an orthogonal check on designs: a high SolvIT score indicates the mutant is less prone to aggregation or inclusion bodies, addressing a common failure mode of stability mutations.

All these metrics are compiled for each design into a single table. The pipeline then merges all metrics with the identity of each sequence and outputs a comprehensive CSV file. This summary contains one row per design and columns for every metric (ESM score, MPNN score, ΔΔG, solubility, RMSD, pLDDT, etc.), including the wild-type as a reference row (with ΔΔG = 0, RMSD = 0, identity = 100%, etc.). The CSV is sorted and easily filterable, allowing researchers to quickly identify the top candidates meeting specific criteria (for example, designs with predicted Tm > +5 °C and RMSD < 1 Å). The webserver version of the pipeline outputs the variants with the highest solubility and thermal stability scores.

#### Implementation and Reproducibility

StabilizeIT is implemented as a Snakemake^13^ workflow, which ensures that all steps are executed in the correct order and that intermediate results are cached. The use of Snakemake enables seamless scaling from local machines to compute clusters, and automatically handles job parallelization and file organization. Each tool in the pipeline (MMseqs2, MPNN, ESM, AlphaFold2, etc.) is encapsulated in a Singularity container image. This containerization guarantees a reproducible environment – all software dependencies (Python libraries, CUDA drivers, etc.) are fixed within the containers. Researchers can thus run the pipeline on any Linux system (including HPC clusters) without worrying about installation conflicts, simply by pulling the provided Singularity images. The pipeline configuration (config.yaml) allows customization of key parameters such as NUM_MPNN_SEQUENCES, SELECT_TOP_K and required database paths. For example, increasing NUM_MPNN_SEQUENCES will sample more designs (improving search coverage at the cost of runtime), whereas lowering SELECT_TOP_K can speed up AlphaFold2 inference if resources are limited. We provide default settings that were found to work well for typical enzyme sizes (300–400 residues), but these can be tuned as needed. In summary, StabilizeIT’s modular Snakemake design and containerization make it a portable and reproducible solution – any researcher can rerun the exact same workflow (recorded with a Git commit hash for provenance]) to reproduce results or extend the pipeline with new modules (e.g. alternative scoring functions or design models).

### Experimental

#### Molecular

The enzyme’s amino acid sequence was reverse-translated to DNA using codon usage optimized according to the E. coli codon frequency table. The resulting DNA fragments were synthesized and cloned into a pET28 vector by Twist Bioscience (CA, USA). The cloned fragments encode an open reading frame (ORF) for the desired enzyme, incorporating an N-terminal 6×His tag for downstream purification.

#### Protein expression

NiCo21(DE3) or BL21(DE3) E. coli competent cells (New England Biolabs, MA) were transformed via the heat shock method^21^. Transformations were performed in a 96-well plate (Cat: 701011, Wuxi-Nest, Jiangsu, China). Following transformation, cells were grown in their respective wells at 37 °C for 16 h in 2xYT medium supplemented with 30 µg/mL kanamycin. Subsequently, cultures were diluted 1:60 into fresh 2xYT medium containing 30 µg/mL kanamycin in deep-well 96-well plates (Labcon, USA, Cat: LC 3909-525) to a final volume of 0.5 mL per well. Cultures were grown at 37 °C with shaking until reaching an optical density of 0.6 _600nm_ and induced using 0.1mM isopropylthio-β-galactoside (IPTG). During induction cells were either grown with shaking at 700 rpm at 25°C for O.N. (**UDP-Glucose 6-Dehydrogenase and β-1**,**4-Galactosyltransferase 2)** or at 16°C for O.N. (**Sucrose Synthase assay)**.

#### Protein isolation and quantification

Cells were pelleted by centrifugation at 4,000 rcf for 15 minutes; subsequently, the pellets were resuspended in 100 µL lysis buffer (BugBuster supplemented with 10 mg/mL lysozyme, a protease inhibitor cocktail, and 0.01 mg/mL Benzonase). The suspension was incubated on ice for 30 minutes and then centrifuged (4,500 g, 4°C, 30 min). The clarified supernatant was mixed in a shaker (800 rpm) with 35µL Ni-NTA Charged MagBeads (GenScript, USA, cat. L00295) for 1 hour at 25 °C. Before incubation, the beads were equilibrated with binding buffer (50mM Tris-HCl pH 8.0, 300mM KCl, 0.1mM ZnCl2, 20 mM imidazole). After binding, the beads were washed twice with 200µL wash buffer (20 mM Tris-HCl pH 8.0, 300 mM KCl, 0.1 mM ZnCl2, 20mM imidazole), followed by a final wash with 200 µL wash buffer containing 50mM imidazole. His-tagged proteins were eluted with 50µL elution buffer containing 50mM Tris-HCl pH 8.0, 100mM KCl, 0.1mM ZnCl2, and 500mM imidazole. Protein concentrations were assessed by SDS–PAGE with Coomassie staining and quantified using the Coomassie Plus Bradford Assay Kit (Thermo Scientific, USA, cat. 23236)

#### Thermal shift assay

Protein thermal stability was assayed using SYPRO Orange dye^22^ (#Cat: 4461146, Applied Biosystems). Measurements were performed with a Bio-Rad C1000 Thermal Cycler and accompanying software. SYPRO Orange was excited at 470nm, and the change in fluorescence emission at 570nm (arbitrary fluorescence units) was monitored over a temperature range from 25°C to 95°C, with an increase of 1°C per minute. Measurement was performed in duplicate; each reaction contained 10µL protein eluate (0.5–1 mg/mL), 2.5 µL pre-diluted SYPRO Orange dye (Thermo Scientific), and 7.5 µL elution buffer. Lysozyme (1mg/mL; Tm 73–74°C) was included as a thermal shift marker. Melting temperatures (Tm) were determined from the inflection points of the fluorescence curves.

#### UDP-Glucose 6-Dehydrogenase assay

Enzyme activity was measured using the reduction of NAD+ to NADH as a proxy for UDP-glucose oxidation. The generation of NADH was monitored at 340nm, where NADH has a molar extinction coefficient of 6,220 M ^-1^cm^-1^. Reactions were performed in standard UV-transparent 96-well plates (UV star) at either 25°C or 45°C. To initiate measurement, 10 µL of purified enzyme was added to a well containing 90µL of assay buffer (20 mM Tris-HCl, pH 8.0; 200mM KCl; 0.1mM DTT; 1mM NAD+; 1 mM UDP-glucose).

#### Sucrose Synthase assay

The conversion of sucrose to UDP-glucose and fructose was monitored using a coupled reaction with UDP-glucose 6-dehydrogenase (UGDH), by tracking the oxidation of UDP-glucose and reduction of NAD+ to NADH at 340nm. Measurements were performed in standard UV-transparent 96-well plates (UV star) using a Tecan infinite M Plex plate reader. To initiate the reaction, 10 µL of purified enzyme was added to 85 µL activity buffer (20mM Tris-HCl pH 8.0, 200mM KCl, 0.1mM DTT, 5mM NAD+, 270mM sucrose, and 16 mM UDP), followed by the addition of 5 µL UGDH (Enzymit UGDH design tyWr4T) final concentration 5 µM. Samples were incubated for 1 hour at 25°C, then absorbance at 340 nm was recorded every 0.5 minutes for a total of 90 minutes at 25°C. Values from the 90 minutes end point were taken for unit calculation. One unit is defined as the amount of the enzyme that catalyzes the conversion of one micromole of UDP-Glucose per minute, using the conversion rate of 2xNADH for one molecule of UDP-glucuronic acid formation at 25°C in an Activity buffer.

#### β-1,4-Galactosyltransferase 2 (GalT2)

The conversion of GlcNAc1P and UDP-glucose to UDP-GlcNAc and glucose-1-phosphate was performed in a standard PCR plate. Five microliters of screened, purified enzyme were added to an activity buffer containing 20mM Tris-HCl (pH 8.0), 100mM KCl, 1 mM MgCl_2_, 3 mM UDP-glucose, and 3mM N-acetylglucosamine-1-phosphate. Reactions were incubated overnight at 25°C. Enzymes were removed prior to HPLC analysis by incubation with 10µL Ni-Mag beads (GenScript) for 20 minutes. Seventy-five microliters of the resulting supernatant were analyzed by HPLC using a HILICpak VN-50 4D column (5 µm, 4.6 × 150 mm). The mobile phase consisted of 70% acetonitrile and 30% 50mM ammonium bicarbonate with 0.3mM EDTA. Runs were performed at 0.75 mL/min at 16°C, using a UV detector set at 280nm, an injection volume of 10µL, and a total run time of 16 minutes. UDP-GlcNAc peaks were quantified using a calibration curve generated with commercial UDP-GlcNAc (#Cat: U4375, Sigma).

## Results

To demonstrate the efficacy of the StabilizeIT pipeline, we applied it to four enzymes of biotechnological interest: **UDP-glucose 6-dehydrogenase (UGDH; Uniprot Id A0A345S0P6), β1**,**4-galactosyltransferase 2 (GalT2; Uniprot Id E8MF11)** and 2 variants of **Sucrose Synthase (SuSy; Uniprot Id Q8DK23, Q820M5)**. These enzymes were chosen for their moderate native stability and importance in various biotechnological applications, making them good candidates for stabilization. For each protein, we provided the wild-type structure (PDB) as input and ran the pipeline with default parameters (100,000 MPNN designs, top 100 selected for modeling).

### Protein Design Evaluation

The StabilizeIT pipeline was applied to the 4 enzymes mentioned above, each cloned into an expression plasmid as detailed in the Experimental Methods section. The resulting designs demonstrate significant improvements across multiple key performance indicators for all four enzymes. Specifically, the majority of the designed variants consistently demonstrated a notable increase in thermal stability, as evidenced by their higher Tm (Figure 2A). This enhanced thermostability is crucial for the practical application and long-term storage of enzymes^23^. Beyond stability, the StabilizeIT pipeline also led to substantial improvements in protein production titers. A significant proportion of the engineered designs exhibited elevated expression titers (Figure 2B). This increase in yield is a critical factor for the cost-effective and scalable production of these enzymes in bioproduction settings. When thermal stability and expression titers were analyzed together—specifically, by examining how many enzymes were both well-expressed and exhibited high melting temperatures (Tm)—the overall success rate remained high. In other words, in all templates, more than half of the designs optimized by our algorithm fell within the intersection of the two sets: enzymes with expression levels higher than the wild type and those with thermal stability exceeding that of the wild type (Figure 2C).

**Figure 1.**
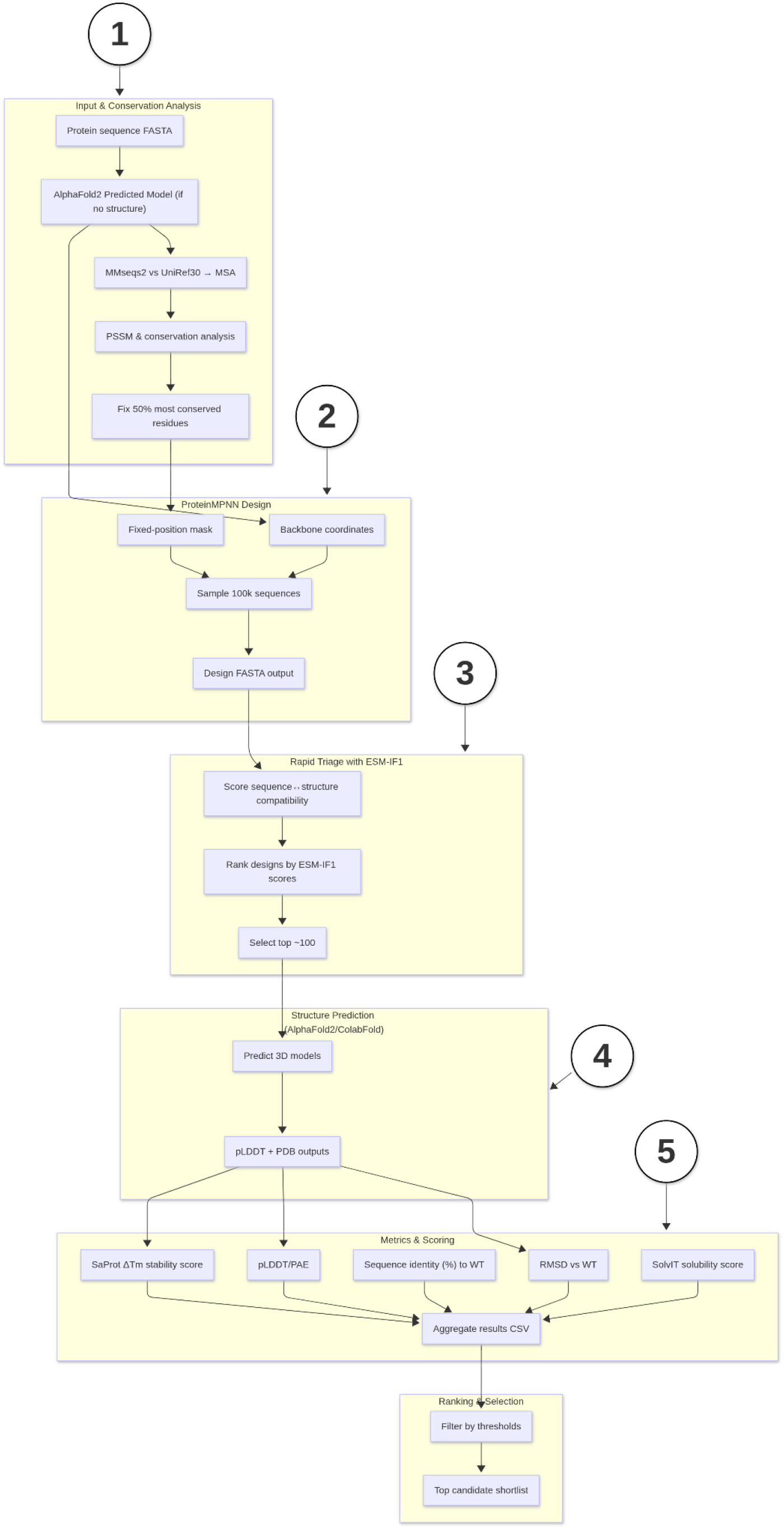
StabilizeIT pipeline flow diagram. Starting from a protein structure or sequence input, the pipeline performs (1) input processing and evolutionary analysis to fix conserved residues, (2) deep sequence design via ProteinMPNN yielding up to N variants, (3) first-pass triaging of designs by ESM likelihood scores to select the top K candidates, (4) structure prediction of top candidates with AlphaFold2/ColabFold, and (5) computation of stability, solubility, and structural metrics for final ranking. The workflow is implemented in Snakemake with Singularity containers for reproducibility.

**Figure 2.**
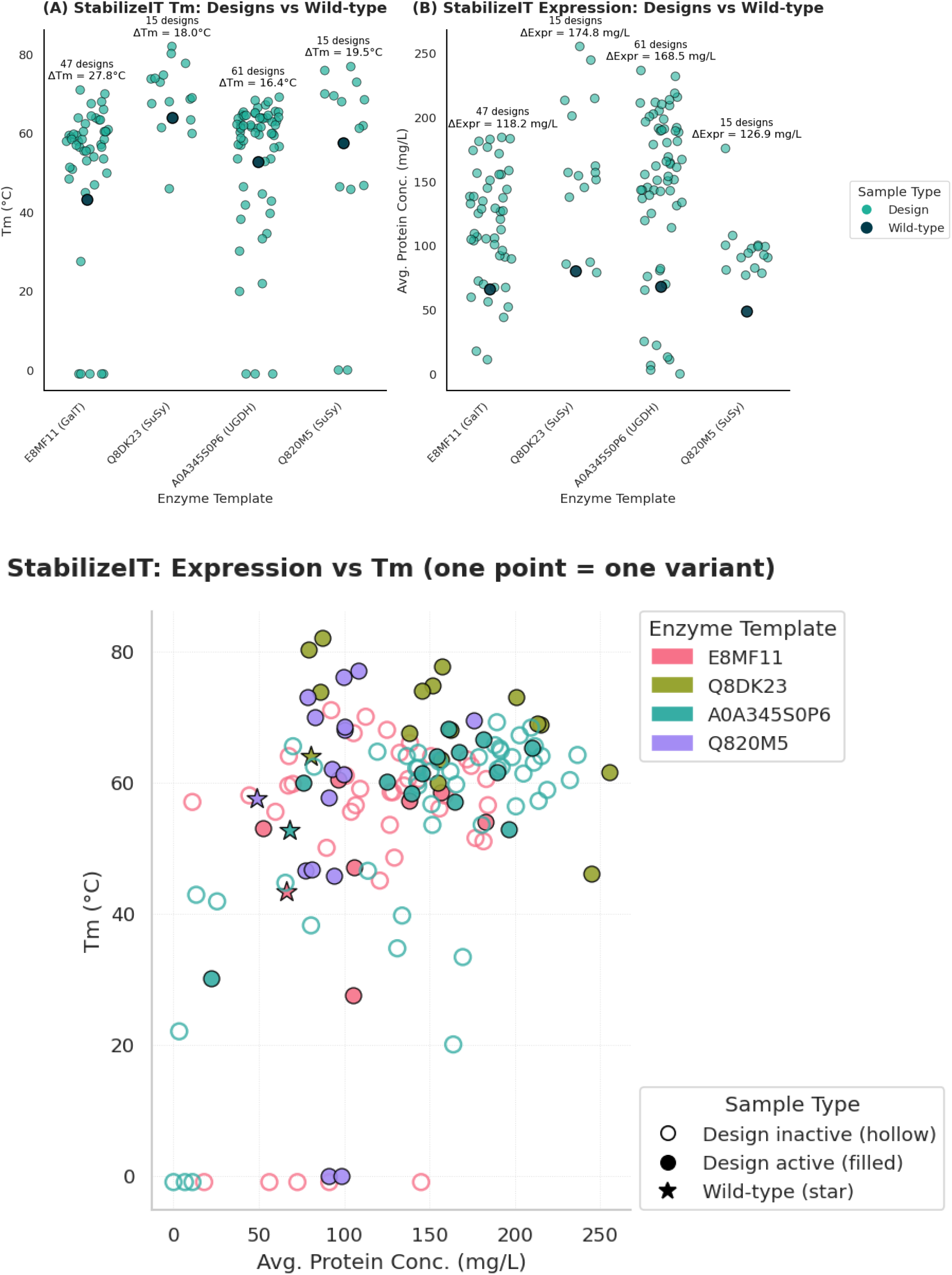
(**A**) melting temperatures (Tm) of engineered enzyme designs vs. their wild-type templates. Comparison of the thermal stability of enzyme variants that were computationally stabilized (“Designs”, in teal) with their corresponding wild-type (WT, dark teal) proteins. Sucrose synthase had insufficient expression and could not be assessed for thermal melting temperature (**B**) Expression titer of engineered enzyme designs vs. their wild-type templates, in mg/L, under identical assay conditions. (**C**) Integrated view. combining panels A and B for every individual variant. Most Design points lie to the upper-right of their WT counterparts, indicating simultaneous gains in thermal stability and soluble expression.

This synergistic enhancement highlights the effectiveness of the StabilizeIT pipeline in optimizing enzyme properties which are crucial for their industrial and research applications.

In addition to expression and thermostability, we assessed whether variants retained catalytic activity using endpoint functional assays. These experiments were designed as qualitative screens, meaning that enzymes were considered active if they achieved at least 10% of the wild-type signal under standardized assay conditions. Across the different enzyme templates, a substantial fraction of designs remained catalytically competent. For GalT (E8MF11), 7 out of 47 designs were active (14.9%). For SuSy (Q8DK23), all 15 tested variants retained activity (100%), and similarly, all 15 variants from SuSy (Q820M5) were active (100%). For UGDH (A0A345S0P6), 13 out of 61 designs demonstrated activity (21.3%). These results indicate that while the fraction of active variants varied by template, the StabilizeIT pipeline consistently produced stabilized sequences that preserved enzymatic function.

The data collectively underscore the pipeline’s ability to generate enzyme variants that are both more robust and more readily produced, thereby accelerating their development and utility.

## Discussion

StabilizeIT provides a modern, unified solution for protein stability design that differs substantially from earlier stabilization platforms such as FireProt, PROSS, and HotSpot Wizard. FireProt^6^ is a web-based pipeline that identifies stabilizing point mutations using a combination of energy calculations (FoldX, Rosetta), evolutionary information, and heuristic filters. It then combines compatible mutations into multi-point variants. While FireProt has produced enzymes with large Tm improvements, its approach is inherently restricted to exploring a narrow mutational space (specific point mutations and their combinations), which limits its ability to capture synergistic or non-intuitive sequence changes. PROSS^5^ also integrates evolutionary information with Rosetta atomistic energy functions. It typically outputs conservative multi-point mutants biased towards consensus sequences (around 10–15 mutations) to ensure function is conserved. The observed increases in thermostability and high expression titers in PROSS designs, however, are largely emergent properties of lowering the protein’s native-state energy and avoiding destabilizing substitutions, rather than the result of explicit optimization for stability, solubility, or expression. Moreover, its reliance on PSSM filtering captures only simple evolutionary likelihoods and fails to account for complex conditional dependencies between mutations. Those observations suggest that while PROSS has achieved notable success across diverse protein optimization tasks, substantial opportunities for improvement remain. HotSpot Wizard^7^ is another protein stabilization framework, which focuses instead on pinpointing structural “hot spots” that strongly affect stability or function. It suggests mutations at these sites using tools like FoldX to avoid destabilizing substitutions. Earlier versions required an experimental structure, while version 3.0 expanded to allow homology-model inputs. While powerful for targeted improvements, its scope is limited to site-specific mutation suggestions rather than broad sequence redesign.

In all three cases, the combination of evolutionary filters and physics-based energy functions represented the state-of-the-art of their time. However, these tools typically generate only a relatively small set of unique variants, and do not leverage advances in deep learning or generative protein modeling. StabilizeIT adopts a fundamentally different approach. By integrating ProteinMPNN with multiple orthogonal predictors, it can generate and triage hundreds of thousands of globally redesigned variants. ProteinMPNN captures complex interdependencies between mutations and backbone structure, enabling redesign of large sequence regions in one step—unlike FireProt or PROSS, which usually keep most of the sequence fixed. This global perspective allows StabilizeIT to escape local optima and identify networks of compensatory mutations across the protein. Safeguards such as conservation masking and active-site RMSD filtering ensure that critical functional regions remain intact. A key differentiator is StabilizeIT’s use of AlphaFold2 as a filtering step, directly testing whether designed sequences are predicted to fold into the correct structure. By requiring high pLDDT scores and low RMSD, the pipeline enforces a stringent virtual screen for foldedness that older methods relying on heuristic energy functions do not provide. Complementary predictors such as ESM inverse folding scores, language-model–based solubility and stability assessments, and SaProt add further robustness and allow for a more fine-grained downstream prioritization of traits (e.g. higher expression titer or thermal stability). Finally, StabilizeIT was built with scalability and modularity in mind. It is implemented with Snakemake and Singularity and can distribute workloads across multiple GPUs, accelerating large-scale design campaigns. Each module is swappable - e.g., AlphaFold2 can be replaced by newer folding models, or alternative generative models like ProGen2 can be plugged in. This open-source, modular design ensures the workflow can evolve with advances in computational protein engineering and is not tied to a particular platform or algorithm. Further metrics can be readily added to assess the feasibility of each of the designs.

